# ReverseScreen.ai: Pharmacophore-Guided Reverse Screening Across the Growing Co-Complex Proteome

**DOI:** 10.64898/2026.09.09.750461

**Authors:** Steven M. Muskal, George Nicola

## Abstract

Biopharmaceutical companies routinely use forward screening to identify potential ligands for argets of biological consequence. Increasingly, they are also using reverse screening to proactively identify downstream off-target liabilities and new repurposing opportunities. Exhaustive reverse docking of one or a multitude of molecules across every characterized site is one approach, but the computational cost is often prohibitive. One molecule against 28,579 receptor sites takes 40.5 hours on twenty CPU cores. We present a method that puts a retrieval step in front of the docking. Every co-crystallized ligand in the Protein Data Bank is indexed by he 3-dimensional pharmacophore fingerprint it presents, together with the UniProt accessions it was solved against, and a query retrieves the sites belonging to its nearest neighbors. Fingerprints are predicted from two-dimensional structures with PharmCast, so a query is fingerprinted in 4 ms and searched against 27,797 indexed ligands in 40 ms. For a query molecule we take its 5 most similar indexed ligands and pool every protein those 5 were crystallized with, which averages 21.1 distinct proteins. Across 3,000 held-out molecules that pool contained the molecule’s own known target 48.8 percent of the time. Docking those 21.1 proteins takes 108 s, against 40.5 hours for the full panel. Pooling the twenty-five most similar ligands instead gives 104.5 proteins and finds the target 60.8 percent of the time. To be indexed, a structure only has to establish which ligand sat in which protein. Holding the index to pocket-grade coordinates had been excluding whole receptor classes. Admitting X-ray at 2.5 Å and cryo-EM at 4.0 Å takes coverage from 3,670 target sites to 28,579. Two 2025 clinical molecules were cross-validated. Orforglipron returned the glucagon-like peptide 1 (GLP-1) receptor first of 27,797 ligands, hrough a non-identical analog at 0.901; daraxonrasib returned the KRAS and cyclophilin A tri-complex fourth, at 0.836. When run in batches the pipeline reverse screens about 96,000 molecules per hour on 1 core, roughly 2.3 million per day, so the retrieval step is practical even with a massive virtual library. The method is available at reversescreen.ai, which allows users the opportunity to screen molecules against the current index, returns the retrieved sites, and docks them individually. The index itself can be downloaded for confidential screening in a local environment.

## 1. Introduction

### 1.1 The reverse question

A screening campaign seeks to identify which ligand in a batch best binds a particular target. The reverse question asks which of many potential targets bind to a particular ligand, and is relevant for a multitude of other questions: what else does a molecule hit, what is behind an unexplained phenotype or a toxicity finding, and what might that compound be repurposed against. It is also the question asked of every molecule a generative method proposes, because a newly-designed structure has no measured target at all.

Chen and Zhi framed the computational form of it with INVDOCK, docking one ligand into a panel of protein cavities and ranking the panel.^1^ TarFisDock^2^ and idTarget^3^ made the approach available as services. The strategy is elegant and its accuracy has been measured: across roughly 470,000 dockings over a panel of 60 targets, the best of 13 docking procedures put the known arget inside the top 10% of the panel for 36% of the 600 query ligands, at a median rank of 11.0 of 60.^4^

### 1.2 Why inverse docking has stayed small

In previous experiments, the panels were relatively small because docking is computationally expensive per site and the resource requirements scale linearly with the panel size. On 20 high-performance CPU cores, AutoDock Vina at “exhaustiveness” setting of 16 takes 5.1 s for a drug-sized ligand. A panel of 60 sites is 5 minutes. The 28,579 sites in this work are 40.5 hours for 1 molecule, which is why published panels have been of the order of hundreds of sites.

The complementary answer is ligand-based and avoids docking. Chemically similar molecules tend to bind the same proteins, so target prediction can proceed from similarity to annotated ligands. Keiser and colleagues related entire pharmacologies this way,^5^ and Daina and Zoete showed that reverse screening of this kind places the correct target first for more than half of a 300,000 molecule set across 2,069 proteins.^6^ It is cheap and works reliably, but the output is a protein name, which stops short of iterative design.

### 1.3 What the crystallographic record already answers

Molecules with similar structure typically behave similarly. A molecule resembling a co-crystallized ligand is likely to bind the same targets that ligand binds. The crystallographic corpus is a rich source of co-complexes and can thus narrow the scope of large scale, multi-target docking exercises. Molecular fingerprints which capture binding features beyond simple 2-dimensional structural representations are well-suited to enhance a virtual screening campaign.

We previously described a method of generating ensemble 3-dimensional pharmacophore fingerprints for QSAR, focused, and primary library design.^7,8^ These fingerprints were limited by the speed of conformer generation and did not scale well. We have recently reported and made available the PharmCast^9^ method which directly predicts a pharmacophoric fingerprint from a SMILES representation, enabling large virtual library exploration and massive compound collection processing. These same PharmCast fingerprints can be used to rapidly identify molecules containing similar pharmacophoric features found in co-crystallized ligands and thus reduce the scope of multi-target docking. And given the scalability of the PharmCast method, this approach can afford massive computational exploration across the co-complex proteome.

The index is a snapshot, not a fixed resource. The Protein Data Bank released 17,608 entries in 2025 and is on course for a similar figure in 2026, about 341 a week, of which about 113 meet the admission rule used here. The qualifying share of new releases has risen over the past decade, from 27.7 percent in 2016 to 36.4 percent in 2025, fluctuating from year to year within that rise (Figure 1). Every result reported below is therefore stated against a dated snapshot.

**Figure 1.**
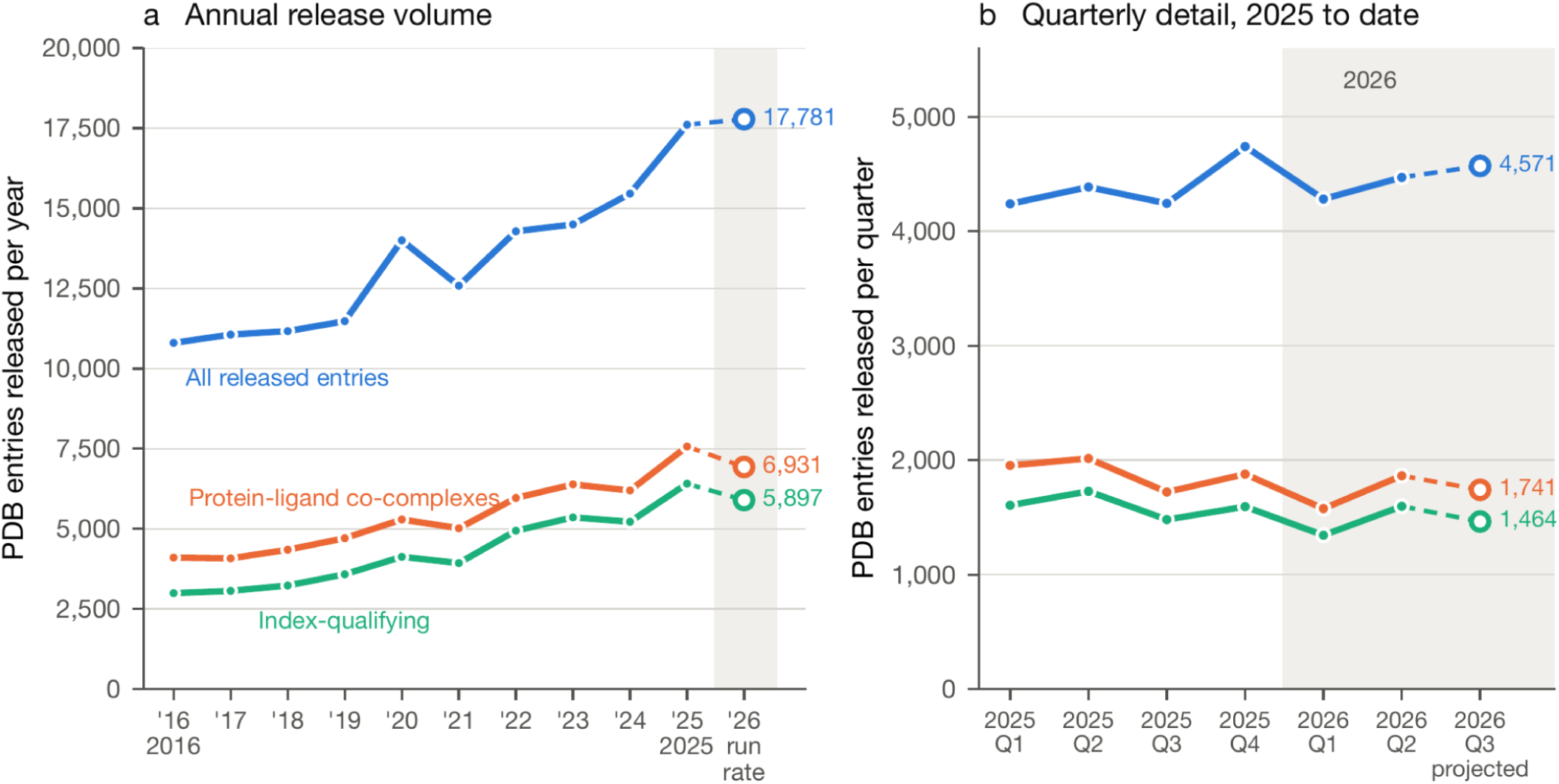
The archive supplies new co-complexes every week. Entries released by the RCSB Protein Data Bank, counted by initial release date and queried on 8 September 2026. (a) Annual release volume. (b) The same three series by quarter since the start of 2025. All released structures are experimental entries. Protein-ligand co-complexes are entries with a polymer entity mapped to a UniProt accession ID that carry a non-polymer entity of formula weight 250 to 900 Da. Index-qualifying entries are that subset meeting the admission rule used here: X-ray at 2.5 Å or better with an R free at or below 0.28, or electron microscopy at 4.0 Å or better. The qualifying share of new releases rose from 27.7% in 2016 to 36.4% in 2025. It fluctuates from year to year within that rise, and the 2026 projection is below 2025 on both filtered series. Open markers are projections from the elapsed fraction of the period: the 2026 annual point is a run rate, and the third quarter of 2026 was 75% elapsed at the query date.

### 1.4 Retrieval before docking

Over the years, we have worked with several biopharmaceutical companies on reverse screening projects; each required months to assemble a panel, prepare receptors, perform the docking and nterpret the results. The panels were small because docking was the bottleneck. The work reported here changes the order: retrieval first, docking second, with the query defining the panel.

The retrieval step is purely ligand based. PharmCast predicts the query’s 10,549-bit pharmacophore fingerprint, which is compared by Tanimoto coefficient with 27,797 indexed igands. The k nearest ligands contribute the UniProt-linked sites from the structures in which hey were solved, and duplicate sites are pooled. The 0.901 similarity in Figure 2 is the comparison between orforglipron and UK1. The receptor enters through the ligand to protein association recorded in PDB entry 6X19.

**Figure 2.**
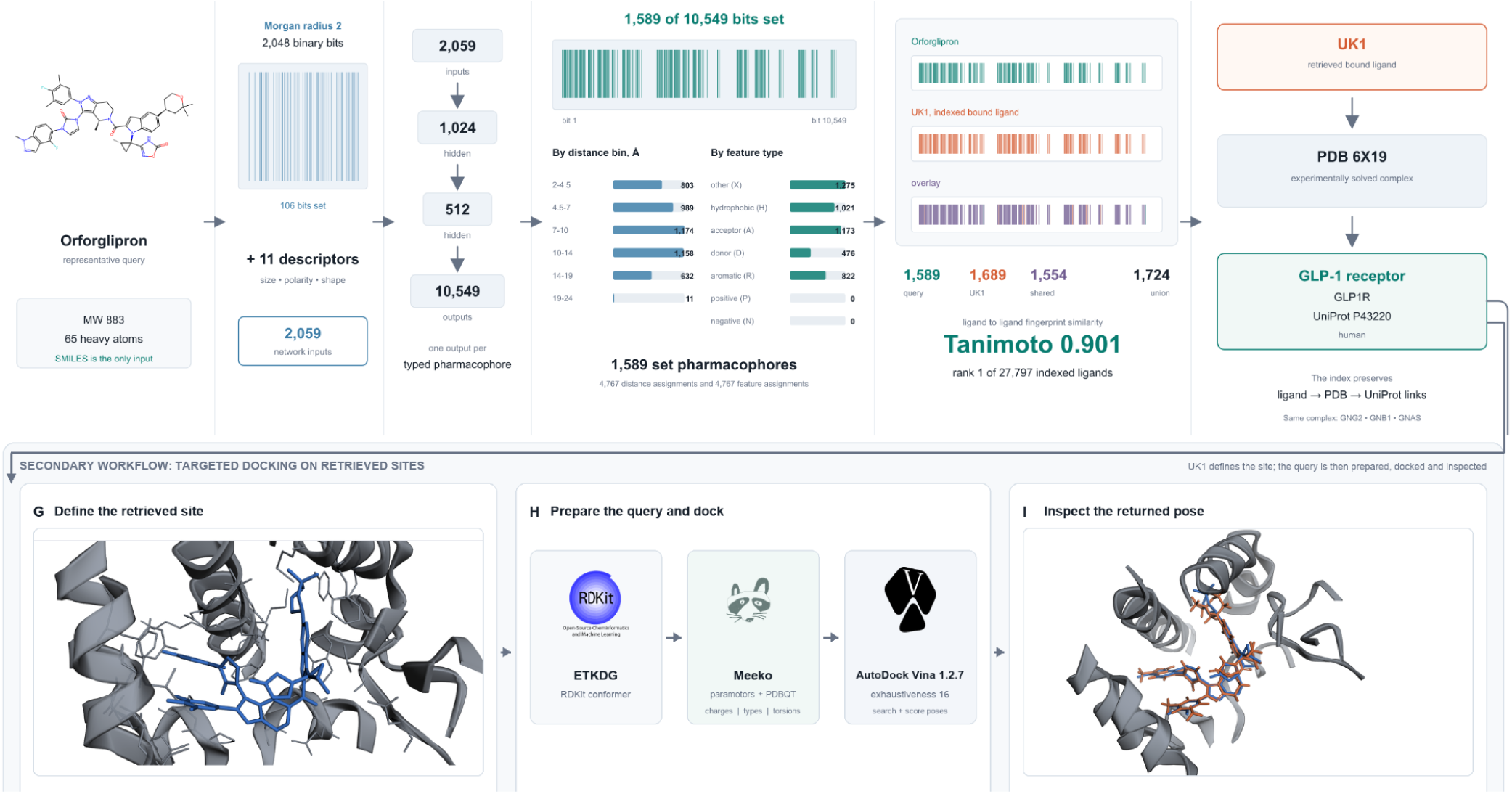
PharmCast retrieval and targeted docking. A query SMILES was encoded as 2,048 Morgan bits and 11 descriptors, then mapped to a 10,549-bit pharmacophore fingerprint. Orforglipron set 1,589 pharmacophores; each contributes 3 distance-bin and 3 feature-type assignments. UK1 shared 1,554 bits Tanimoto 0.901) and linked the query to GLP-1R through PDB 6X19. Panels G-I show the 6X19 drill-down: deposited UK1 defines the search box, expanded by 4-5 Å; RDKit’s Experimental-Torsion Knowledge Distance Geometry (ETKDG) method generated the orforglipron conformer; Meeko was used to prepare it, and AutoDock Vina 1.2.7 was used to dock it at exhaustiveness 16. The orange pose scored −15.60 kcal/mol; UK1 is displayed in blue. One pose was returned in 366.8 s. A separate redocking control placed the top pose 1.02 Å from the deposited 6XOX pose. Interactive pose is available at: https://reversescreen.ai/pose.html?id=6X19-u8w6ct0B

Docking is the subsequent drill-down step on that candidate pool. For each site, the indexed igand locates the pocket and defines a search box from its envelope plus 4 to 5 Å. The query is hen prepared and docked into the retained receptor structure. During the retrieval step, an optimal site is selected. The docking step then provides poses that can be inspected and scored. The complete sequence is shown in Figure 2.

We provide recall results against docking cost on 3,000 held-out queries, redocking accuracy on 38 complexes, and two 2025 clinical molecules run end to end with their own structures left out.

## 2. Methods

### 2.1 Lowered bar on a “good” structure

A method that compares one binding site to another has to superimpose every atom that lines the pocket. Pseudocenter positions, interaction geometry and surface shape are all derived from coordinates, and an error of half an angstrom in a side chain moves a feature into a different distance bin. Work of that kind is properly restricted to well-refined crystallography,^10,11^ and the rule commonly used elsewhere is X-ray diffraction at 1.8 Å or better with a free R factor at or below 0.23.

Retrieval asks a different question of the same structure repository. It does not compare pockets and it does not use the receptor at all. It needs the identity of the ligand, the fingerprint that igand presents, and the accession ID of the protein it was solved against. A 3.1 Å cryo-EM reconstruction establishes that association as securely as a 1.2 Å crystal structure does, even hough it would be ineffective at describing the pocket.

Applying the stricter rule to both questions loses coverage silently. The loss falls unevenly. It falls on the receptor and large-complex classes whose structures are cryo-EM. The glucagon-like peptide 1 receptor is the clearest case in this work: it carries 31 entries holding a drug-like ligand, and none of them satisfies the 1.8 Å rule, so under that rule the receptor is absent from the panel and no amount of searching can return it. The retrieval index therefore admits X-ray entries at 2.5 Å or better with a free R factor at or below 0.28, and cryo-EM entries at 4.0 Å or better, in both cases carrying a non-polymer component of 250 to 900 Da.

### 2.2 Building the index

Entries were enumerated through the RCSB search service on September 8, 2026 under the two rules above, giving 61,935 X-ray entries and 12,620 cryo-EM entries. The cryo-EM scan contributes 15,281 target sites and the X-ray scan 14,437, overlapping where an accession has been solved by both methods, so just under half of what the index can retrieve against, 14,142 of 28,579 sites, would be absent under an X-ray rule alone. For each entry, every non-polymer component in the mass window was paired with every UniProt accession carried by the entry’s polymer entities. A ligand in a structure containing two chains was recorded against both, because the purpose of the index was to propose sites for docking and the docking stage resolves which chain the ligand can reach. The Protein Data Bank releases new entries once a week, on Wednesday. The index behind reversescreen.ai is scheduled to rebuild each Sunday, so a release becomes searchable within five days. Every screening result reports the snapshot it was produced against, and the interactive website at reversescreen.ai returns the current snapshot, its build time, and the admission rule at reversescreen.ai/api/index-info. The website is configured for nteractive use, and its defaults are not the settings benchmarked here.

Component definitions were read from the chemical component dictionary. SMILES strings attached to entries were not used, so each molecule carries its deposited atom names and bond orders. A component built from a flat SMILES loses the bond order assignment that distinguishes an aromatic ring from a saturated one, and the pharmacophore fingerprint is sensitive to exactly that distinction: a pyrazole and a pyrazolidine present different features from the same skeleton.

Of the components paired to an accession ID, 237 were dropped because their definition would not build a valid molecule or because their mass fell outside the window once hydrogens were assigned. The index holds 27,797 components covering 28,579 target sites, against 3,670 sites under the stricter rule, an increase of 7.8 fold. Every count in this manuscript was taken during the September 8, 2026 snapshot. As the archive grows with each weekly release, a build taken after further releases returns larger totals and the recall reported here becomes a lower bound for it.

### 2.3 Fingerprints

Each molecule is described by a 3-dimensional pharmacophore fingerprint. A bit is one 3-point pharmacophore, a triple of typed features with the 3 inter-feature distances binned, and the molecule sets that bit when any accessible conformation can present the triangle.^7,8^ The computational cost is related to its conformation. Generating a triangle requires a conformation that presents it, so the reference calculation generates an ensemble and unions the bits across it. In such a pipeline conformer generation consumes 2.82 s of the 2.86 s spent per molecule.^9^ At 27,797 components an index build would take 22 hours, and at virtual library scale the calculation is prohibitive.

In our method, the fingerprint is predicted instead of computed. PharmCast is a feedforward neural network that returns all 10,549 bits of the ensemble fingerprint from a 2-dimensional structure, reproducing the reference calculation at a Pearson correlation of 0.980 with held-out screening compounds.^9^ Every indexed component and every query is predicted the same way, so the two sides of a comparison are never a mixture of a predicted and a computed fingerprint. The complete index is fingerprinted in 29 s.

### 2.4 Retrieval and its cost

Fingerprints are packed to 1,319-byte vectors and a query is compared against the whole index by Tanimoto coefficient, computed from population counts over a bitwise AND. The search is an exhaustive scan, without an approximate index, clustering, or pruning. This keeps it exact and removes any parameter that could be tuned to the benchmark. The sites carried by the *k* nearest components form the candidate pool, and where several of the nearest components carry the same site, that site enters the pool once.

### 2.5 Held-out evaluation

3,000 components were drawn at random from the index and used as queries. Before each query is searched, every component sharing its InChIKey connectivity layer is removed, so a molecule can never retrieve a site through itself, through a stereoisomer of itself, or through a redeposition of itself under a different component code. The connectivity layer is used deliberately: a query and a deposited stereoisomer of that query are the same retrieval, and counting one as a success would inflate the result.

The ground truth for a query is the set of PDBs its own component is recorded against, and a query counts as recalled when the pool contains any of them. Recall is the criterion because the pool is what the docking stage receives. A site ranked second and a site ranked twentieth are comparable if both are successfully docked. A site outside the pool is eliminated from the pipeline.

### 2.6 Docking

AutoDock Vina 1.2.7^12,13^ was used for docking, and ligands were prepared by Meeko with input conformers generated using ETKDG.^14^ The search box was centered on the retrieved site’s own co-crystal ligand and sized to that ligand’s envelope plus 4-5 Å. An oversized box was not a safe default: padding a large reference ligand by 8 Å gave boxes near 23,000 Å³, and at that volume the search returned surface poses for both case molecules.

Receptors retained cofactors and metals and dropped only waters and the ligand being placed. Deleting every heteroatom removed the guanosine diphosphate and magnesium that sit inside the KRAS site, and the search then filled the cavity the cofactor should occupy.

The redocking panel was 38 complexes generated from the X-ray structures that meet the site rule, one per UniProt accession, with ligands between 250 and 600 Da. Every pose Vina wrote was also rescored with the Vinardo function through the same binary, and the pool was re-ranked by Vinardo alone and by the mean of the two ranks.

Redocking discarded the deposited pose entirely, generated a fresh conformer, and supplied only he box, so no crystal geometry other than the location of the site reaches the docker. Redocking establishes that the search and scoring can place a ligand in a pocket the ligand is known to occupy. It does not establish the accuracy of a pose at a retrieved site.

## 3. Results

### 3.1 The pipeline and its cost

**Figure 3.**
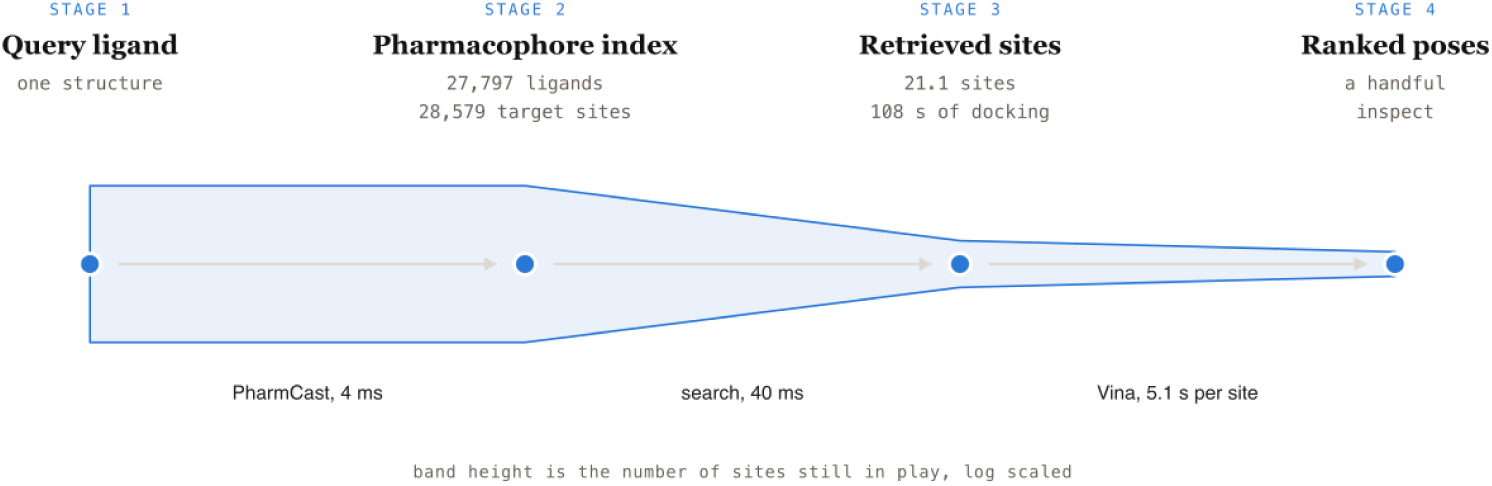
The reverse screen. Band height is the number of receptor sites still relevant at each stage, log scaled. Retrieval removes 99.9% of the panel in 44 ms, which was what made the docking stage affordable.

### 3.2 Coverage

The retrieval rule admits 28,579 target sites against 3,670 under the site fingerprint rule, an ncrease of 7.8 fold. The gain was not uniform across target classes: it was concentrated in receptors and large complexes whose structures are cryo-EM.

**Figure 4.**
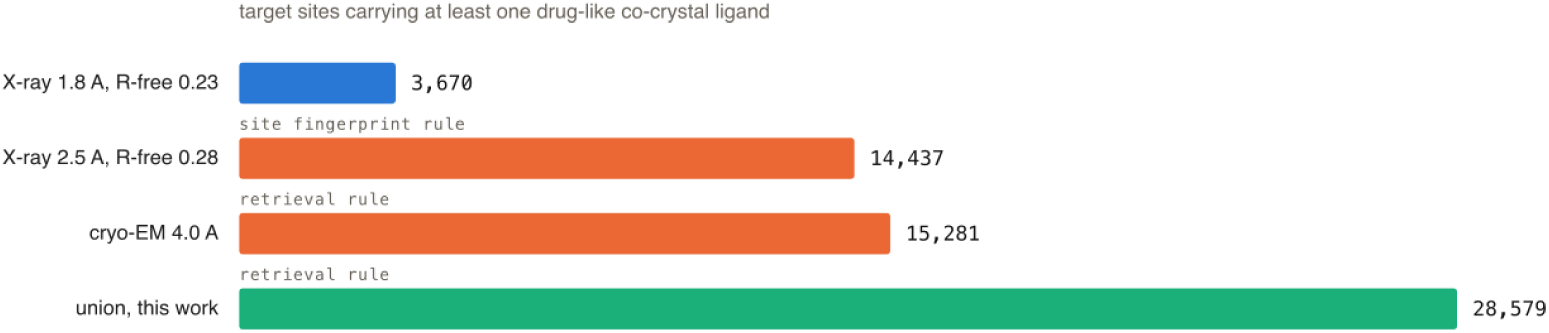
Target sites admitted by each structure rule. The site fingerprint rule and the retrieval rule target different questions of the same repository.

The index was not uniform. Embedding all 27,797 ligand fingerprints in three dimensions separates them into regions that correspond to target class, and both case molecules sit inside the region holding the ligands that retrieved their targets. The embedding was a navigational aid rather than validation. Recall was measured and described in section 3.4.

**Figure 5.**
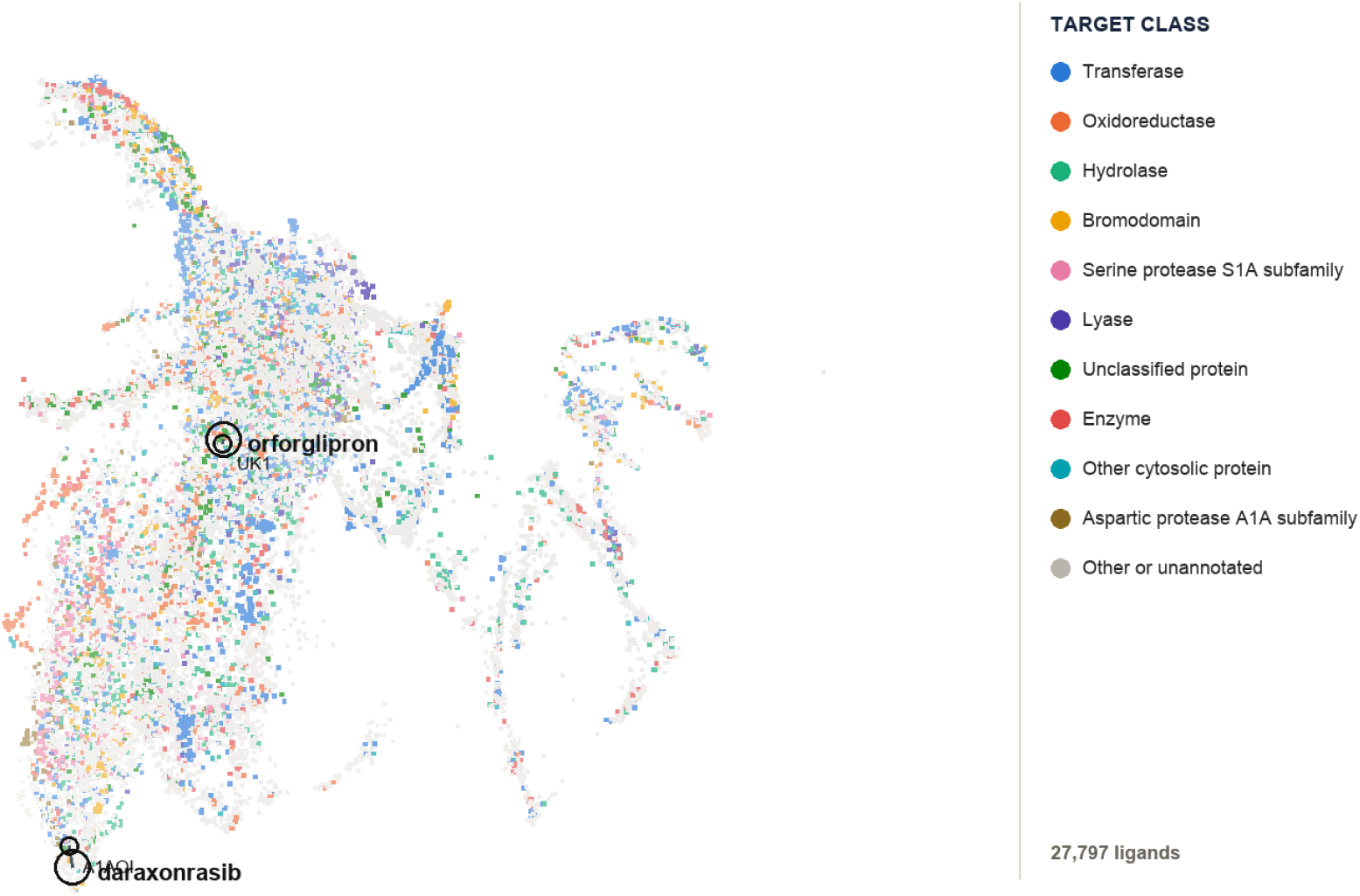
The retrieval index in three dimensions. One point per indexed co-crystal ligand, embedded by UMAP directly from the full 10,549-bit fingerprints under Jaccard distance, which the Tanimoto coefficient etrieval uses on binary vectors. Nothing was reduced or compressed first, so the distances drawn here are the distances the search computes. Points are colored by the target class the ligand was solved against. Orforglipron and daraxonrasib are ringed and labeled, each joined to the indexed ligand that carried its true arget.

### 3.3 Throughput

A single query was fingerprinted in 4 ms and searched in 40 ms. Screened in batches, they average 37.3 ms per molecule, or about 96,000 molecules per hour on one core and roughly 2.3 million per day. Against a 27,797 ligand index it is approximately 2.68 billion pairwise fingerprint comparisons per hour.

At a batch of 1,000 the prediction of 1,000 fingerprints takes 0.4 s and the search takes 36.9 s, so he search, rather than the fingerprinting, becomes the bottleneck. The scan was kept exhaustive because it was exact and has no parameter to tune, and at 99 percent of the batch time it was the step that would result in an approximate nearest neighbor structure if the index grew by another order of magnitude. At this index size it was already fast enough that docking dominated the pipeline by three orders of magnitude.

### 3.4 Recall against docking cost

The procedure was the same at every setting and only the number of neighbors changes. It takes he query’s *k* most similar indexed ligands, pools every protein those ligands were crystallized against, and asks whether the query’s own target was in the pool.

Recall increases with the pool, which grows faster than the recall does. Five neighbors were the operating point we recommend for a library: 21.1 proteins, 108 s of docking, and the true target present for 0.488 of queries. Table 1 gives the full range and Figure 6 carries the same measurement past it. Recall was close to linear in the logarithm of the pool, so each tenfold ncrease in docking buys about 18 more recall points. Pooling the whole index gives 0.972, because for 2.8 percent of queries no other indexed ligand touches the true site at any depth. The price of a recall point rises with the pool, from 0.6 minutes of docking between 5 and 25 neighbors to 53.4 minutes between 1,600 and 3,200. A single molecule run on its own was not held to the library setting: 400 neighbors reached 0.815 for 1.42 hours of docking, which was affordable once but prohibitive when run 100,000 times.

**Figure 6.**
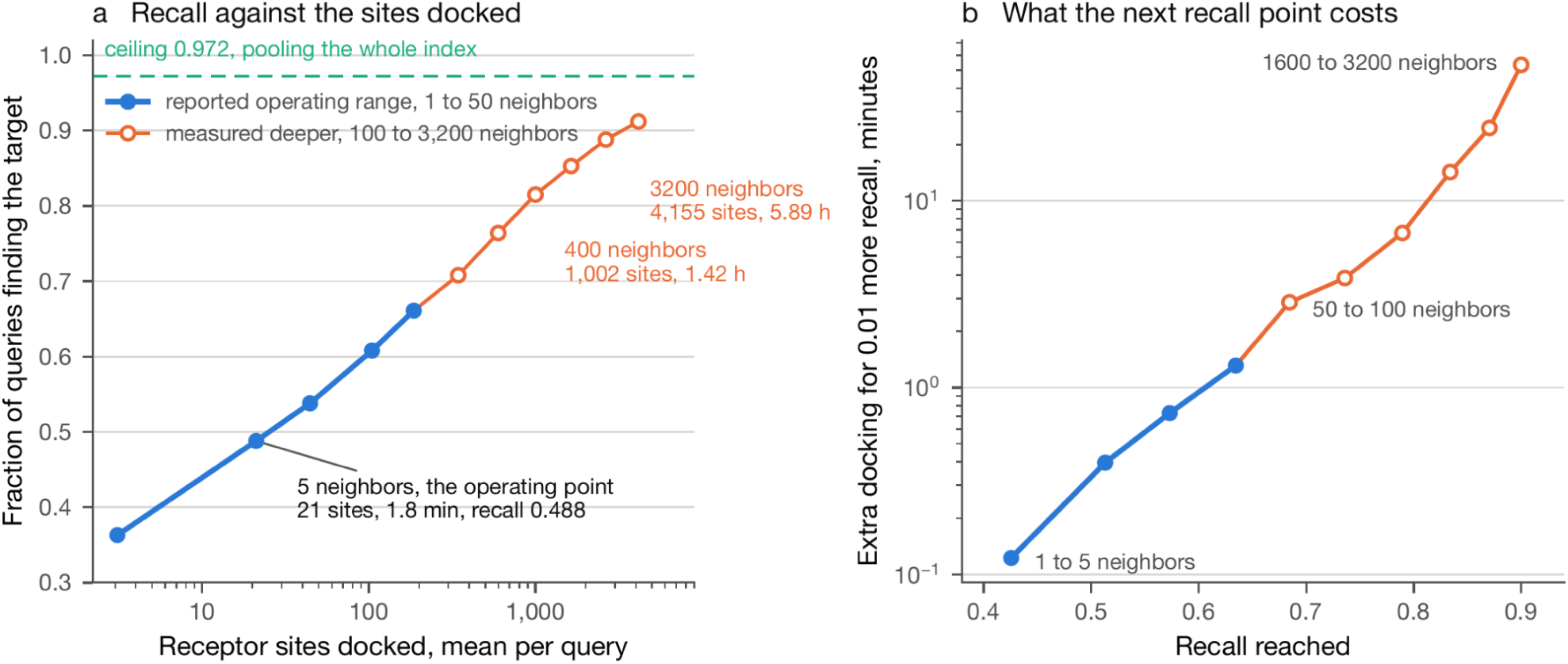
Recall against the sites docked, and the price of buying more of it. (a) Recall over 3,000 held-out queries against the mean number of receptor sites in the pool, logarithmic. Filled markers are the rows given n Table 1, drawn to 3,200 neighbors. The dashed line was the ceiling, 0.972, reached by pooling the whole ndex. (b) The extra docking afforded 0.01 more recall, by band. Every point was measured. Recall at a pool depth was the fraction of queries whose own target first appears at or before that depth, so a single pass over he queries gives recall at every depth.

**Table 1.** Each row pools the proteins bound by the query’s *k* most similar indexed ligands and reports how often the query’s own target was among them, over 3,000 held-out molecules. Docking time was the pool size at 5.1 s per protein, against 40.5 hours for all 28,579.

| Nearest ligands pooled | Fraction finding the target | Mean proteins in the pool | Docking time | Against all 28,579 |
| --- | --- | --- | --- | --- |
| 1 | 0.363 | 3.1 | 16 s | 9214 |
| 5 | 0.488 | 21.1 | 108 s | 1353 |
| 10 | 0.538 | 44.4 | 3.8 min | 643 |
| 25 | 0.608 | 104.5 | 8.9 min | 274 |
| 50 | 0.661 | 186.3 | 15.8 min | 153 |
| 100 | 0.708 | 344.3 | 29.3 min | 83 |
| 200 | 0.764 | 598.5 | 50.9 min | 48 |
| 400 | 0.815 | 1,001.5 | 1.42 h | 29 |
| 800 | 0.853 | 1,638.1 | 2.32 h | 17 |
| 1600 | 0.888 | 2,647.9 | 3.75 h | 11 |
| 3200 | 0.912 | 4,155.2 | 5.89 h | 7 |
| all 27,797 | 0.972 | 28,578.5 | 40.5 h | 1 |

### 3.5 Redocking

The docking stage was validated by redocking: the deposited pose was discarded, a fresh conformer was generated, and the ligand was docked back into its own receptor. A pose counts as correct at 2.00 Å from the deposited coordinates. Over 38 complexes the search generates a correct pose in 27 of them. The scoring function puts that pose first in 14 and within the first hree in 23. On 3DJX the correct pose was present at 1.97 Å and scored sixth.

**Table 2.** Redocking 38 complexes at exhaustiveness 16 on 20 cores, ordered by the rank at which the first correct pose appears. RMSD in angstroms against the deposited pose, correct at 2.00 Å. A rank of none means no pose in the pool reached 2.00 Å.

| Complex | Ligand | Poses | Top pose RMSD | Best pose RMSD | Rank of first correct pose | Seconds |
| --- | --- | --- | --- | --- | --- | --- |
| 9JVI | A1EDG | 7 | 0.37 | 0.37 | 1 | 2.9 |
| 7ZIH | JCR | 9 | 0.42 | 0.42 | 1 | 5.4 |
| 5MZI | FYK | 6 | 0.46 | 0.46 | 1 | 4.0 |
| 4MP7 | PFT | 9 | 0.68 | 0.68 | 1 | 2.2 |
| 5G4N | O83 | 9 | 0.70 | 0.70 | 1 | 2.2 |
| 6MGC | C5P | 9 | 0.77 | 0.77 | 1 | 6.3 |
| 5CTX | 55G | 9 | 0.77 | 0.77 | 1 | 3.5 |
| 3MHJ | M3F | 4 | 0.84 | 0.84 | 1 | 1.8 |
| 6S3A | SFX | 9 | 0.93 | 0.93 | 1 | 4.4 |
| 2QXW | LDT | 9 | 0.96 | 0.96 | 1 | 5.2 |
| 6XOX | V6G | 9 | 1.02 | 1.02 | 1 | 101.8 |
| 1US0 | LDT | 9 | 1.11 | 1.11 | 1 | 5.5 |
| 4JU9 | TZD | 9 | 1.39 | 1.39 | 1 | 11.1 |
| 9HDW | U5P | 9 | 1.74 | 1.53 | 1 | 4.5 |
| 6UYH | HFG | 7 | 2.05 | 0.41 | 2 | 4.3 |
| 9AX6 | A1AHB | 7 | 3.67 | 0.87 | 2 | 217.0 |
| 7LYE | VRD | 9 | 4.10 | 0.75 | 2 | 7.2 |
| 9U7G | A1EOH | 9 | 6.89 | 1.66 | 2 | 11.4 |
| 7ZXZ | K9R | 9 | 3.36 | 1.01 | 3 | 14.9 |
| 9NWX | A1B66 | 9 | 4.22 | 1.08 | 3 | 5.7 |
| 2F6V | SK2 | 9 | 5.85 | 1.83 | 3 | 6.8 |
| 4LBS | 4O8 | 9 | 7.37 | 0.84 | 3 | 5.6 |
| 9Y8R | A1CTP | 9 | 7.78 | 0.92 | 3 | 5.0 |
| 4QFH | G6P | 9 | 5.47 | 1.24 | 4 | 3.9 |
| 9TMX | FB8 | 9 | 6.87 | 1.53 | 4 | 4.1 |
| 6TGU | N92 | 9 | 2.26 | 0.50 | 5 | 4.2 |
| 3DJX | C5P | 9 | 4.83 | 1.97 | 6 | 5.5 |
| 8J7I | 1SM | 9 | 2.15 | 2.15 | none | 4.4 |
| 8OKG | VQH | 9 | 2.26 | 2.26 | none | 3.1 |
| 5LW3 | B3P | 9 | 3.47 | 3.17 | none | 10.5 |
| 5QBH | D8M | 9 | 4.37 | 4.36 | none | 4.8 |
| 4X7I | 3YS | 9 | 4.50 | 4.50 | none | 3.9 |
| 7G1G | WO0 | 9 | 5.20 | 2.60 | none | 2.3 |
| 6T2D | M9E | 9 | 5.84 | 4.58 | none | 4.2 |
| 3TP6 | THP | 9 | 6.00 | 2.66 | none | 7.3 |
| 4TQC | M7G | 9 | 6.04 | 5.29 | none | 10.9 |
| 5MME | 8Q6 | 9 | 7.21 | 3.05 | none | 5.6 |
| 1H01 | FAL | 9 | 9.94 | 9.41 | none | 9.9 |

**Figure 7.**
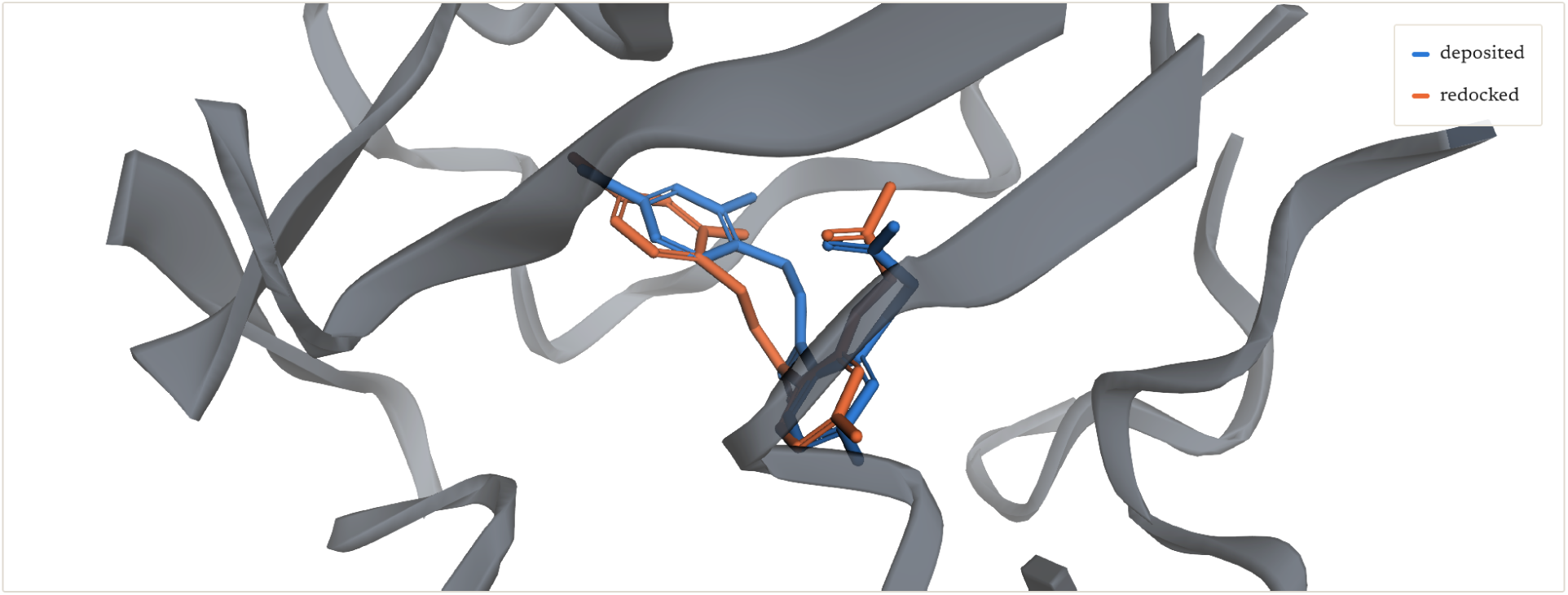
Human aldose reductase 1US0 at 0.66 Å^15^ with the deposited IDD594 pose and the top-scored edocked pose, 1.11 Å apart.

### 3.6 Orforglipron

Orforglipron is an oral non-peptide agonist of the GLP-1 receptor, 883.0 Da and 65 heavy atoms. It is deposited as component V6G in 6XOX,^16^ which was removed from the index for this run. Fingerprinting and search took 43.5 ms together.

The nearest indexed ligand was UK1 at 0.901, a non-identical analog from 6X19,^17^ and it carries he receptor. The true target was returned at rank 1 of 27,797. A pool of twenty-five neighbors holds 48 sites.

**Table 3.** Nearest indexed ligands for orforglipron with the molecule itself withheld. Rows carrying the true arget are marked.

| Component | Tanimoto | Target sites |
| --- | --- | --- |
| **UK1** | **0.901** | **P43220, P59768, P62873** |
| E40 | 0.803 | P14780 |
| IHM | 0.796 | Q08499 |
| D84 | 0.790 | P01375 |
| 7TM | 0.789 | Q02293, Q04631 |
| IC7 | 0.787 | P03372, Q15788 |
| 5IX | 0.786 | A0A8V0Z8P0, P63043, P81947 |
| A1AZI | 0.786 | P62942 |

**Figure 8.**
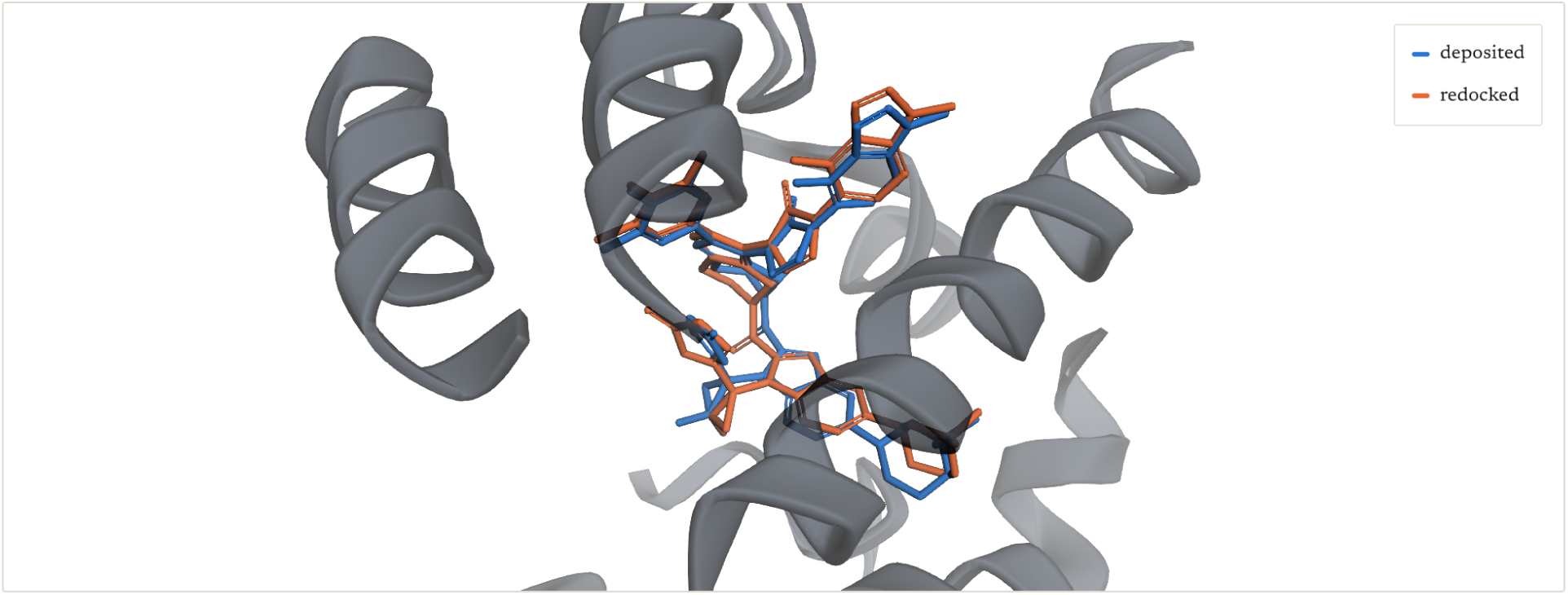
Orforglipron redocked into its own deposited entry 6XOX, shown against the deposited pose. Top pose 1.02 Å, Vina score −14.51 kcal/mol. The docking into the retrieved site, 6X19, is in Figure 2.

### 3.7 Daraxonrasib

Daraxonrasib is a macrocyclic RAS(ON) multi-selective inhibitor, 811.1 Da and 58 heavy atoms, hat acts through a tri-complex with cyclophilin A. It is deposited as component A1AHB and was withheld for this run.

The nearest indexed ligand had a similarity of 0.863 and did not carry the target. The first ligand hat does is A1AOI at 0.836, rank 4 of 27,797, and a pool of twenty-five neighbors holds 50 sites. A macrocycle with no close analog in the record was therefore recovered by the same procedure, at a lower similarity and a slightly deeper pool.

**Table 4.** Nearest indexed ligands for daraxonrasib with the molecule itself withheld.

| Component | Tanimoto | Target sites |
| --- | --- | --- |
| 3N6 | 0.863 | A0ACD6BAW2, O60894, Q16602 |
| A1BB4 | 0.843 | P40337, P51532, Q15369 |
| GGI | 0.842 | P00747 |
| **A1AOI** | **0.836** | **P01116, P62937** |
| KD7 | 0.832 | Q9Y5Y6 |
| 436 | 0.815 | Q13490 |
| GT6 | 0.815 | P98170 |
| OPT | 0.810 | P07711 |

**Figure 9.**
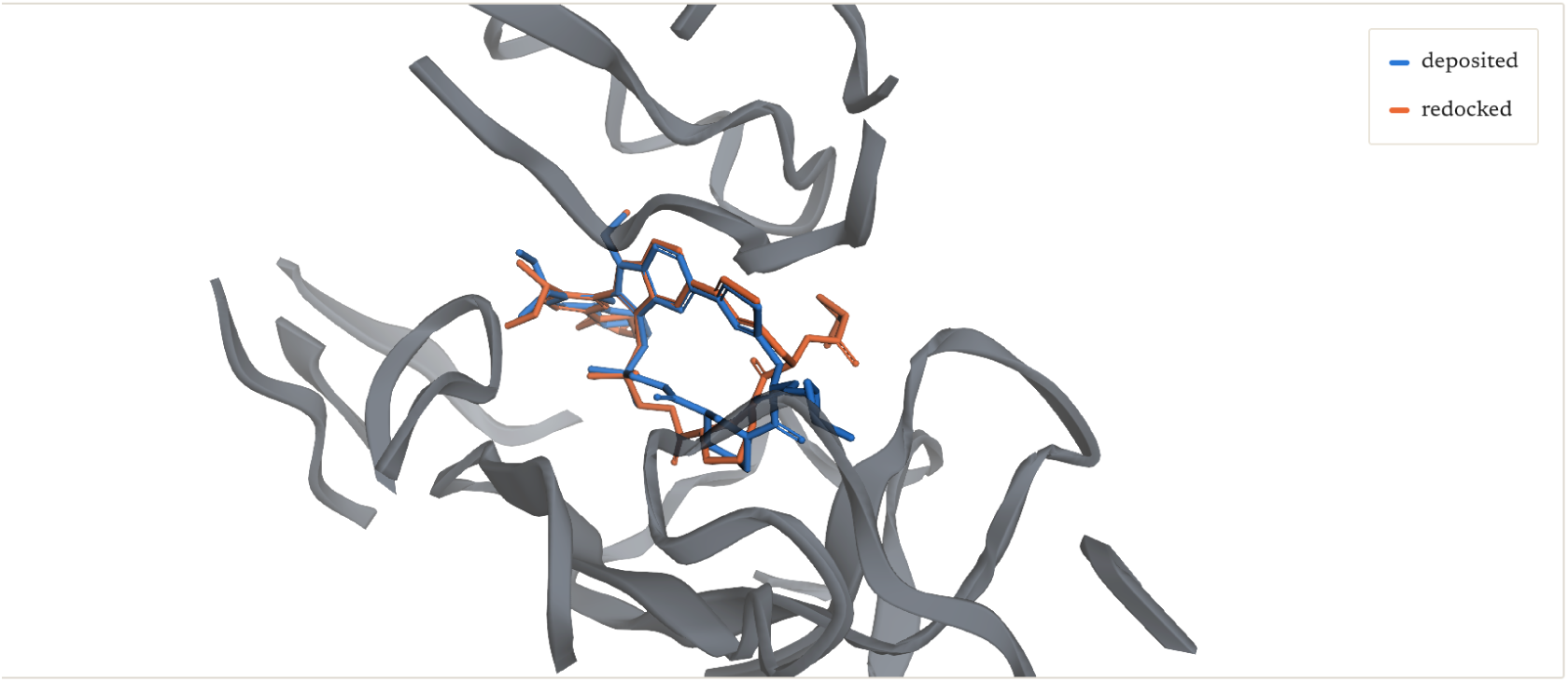
Daraxonrasib redocked into its own deposited entry 9AX6 at 1.65 Å, the KRAS and cyclophilin A ri-complex, shown against the deposited pose.^18^ Top pose 3.67 Å, Vina score −10.59 kcal/mol.

## 4. Discussion

Retrieval changes the scheduling problem. Reverse screening projects we have run with collaborators were measured in months. The panel size was set by how much docking fit in the schedule, not by how much of the proteome was characterized. With retrieval in front, one molecule against the whole characterized proteome took a few minutes and a library of 100,000 molecules took an hour of retrieval plus the docking budget the campaign chose to spend on the pools.

Historically, reverse screening has been limited by the panel it can afford to dock. Putting pharmacophore retrieval in front of the docking changes that panel from a fixed list of dozens to he whole structurally characterized proteome, because each molecule is able to select its own. A query takes 44 ms to place and returns a pool that takes 1-9 minutes to dock, against the exhaustive alternative.

The two admission rules matter as much as the retrieval. The GLP-1 receptor is represented only by cryo-EM entries, so the stricter rule excluded it from the panel altogether. Separating the rules made it reachable, and retrieval then returned it at rank 1.

Recall is set by what has been crystallized. A pool that does not contain the true site cannot be repaired downstream, so the index is the object to grow, and new depositions are what extend it. The retrieval itself was already fast enough to be free relative to a single docking, and its cost does not change with the depth of the pool, because the scan ranks the whole index whatever depth was taken from it. The operating point was therefore set by the docking budget, and it was not one number. A single molecule can afford the pool that reaches 0.815, which was 1.42 hours on 20 cores. A library of 100,000 cannot, so 5 neighbors was the setting for a library and 400 the setting for a single molecule.

The remaining error lies principally in scoring. Vina generates a correct pose in 27 of 38 redockings and ranks it first in 14, so the pool holds a correct pose far more often than the top score names one. Rescoring every pose with Vinardo puts the correct pose first in 12 and within he first three in 24. Ranking by the mean of the two functions gives 14 and 24. Neither improves on Vina at the top of the list. Carrying the first 3 poses forward instead of the first raises the correct pose from 14 to 23 of 27. That was the operating point to use. Three poses per site costs nothing extra, because the pool was already small.

### Extending the index beyond the experimental archive

The index was built from experimental co-complexes because a co-crystal structure is a measurement of where a ligand sits, whereas a modeled complex is a hypothesis about the same thing. Several large collections of modeled complexes now exist. AlphaFill^22^ transplants ligands from solved structures into predicted models by superposition of homologues, and holds about 17 million transplanted ligands across more than 840,000 proteins under a license permitting both academic and commercial reuse. Co-folding methods including Boltz, Protenix and OpenFold generate complexes directly and are openly licensed. Docked pose collections such as CrossDocked2020 place ligands into pockets computationally at a scale of tens of millions of poses.

We examined AlphaFill in detail because it is the largest of these. Its ligand geometries are transplanted from crystal structures and carry a different, more favorable error profile than generated poses. Most of its contents are not the kind of molecule a reverse screen is tasked with. Half of all transplanted ligands are bare metal ions and a fifth fall below the mass floor used in he current study. Of the remainder, close to half are natural cofactors, and most of the remainder are lipids, detergents and buffers. Requiring a component in the mass window used throughout this work, with at least 2 rings, at most 10 rotatable bonds and at least 1 nitrogen, and excluding cofactors and crystallization additives, retains under 6% of the transplanted ligands, and about 1 protein in 8 once the resource’s own high confidence criteria on local backbone deviation and steric clash are also applied (Figure 10).

**Figure 10.**
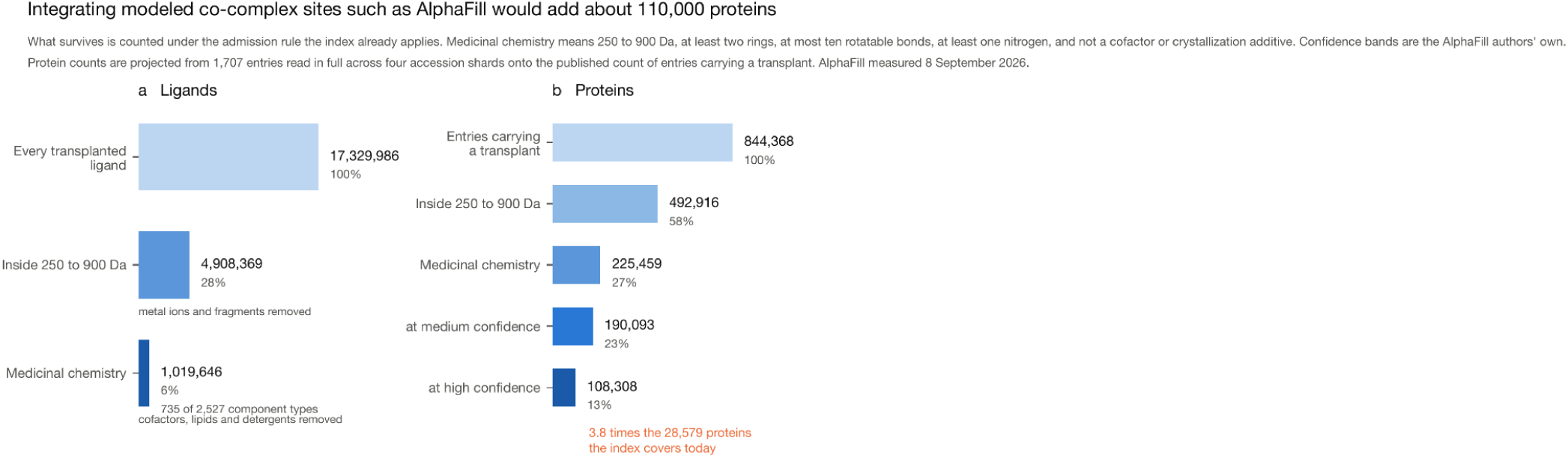
Integrating modeled co-complex sites such as AlphaFill would add about 110,000 proteins. AlphaFill transplants ligands from solved structures into predicted models by superposition of homologues, measured 8 September 2026. (a) Transplanted ligand instances. Half are bare metal ions and a fifth fall below the 250 Da floor used here; of what remains, close to half are natural cofactors, and most of the rest are lipids, detergents and buffers. Requiring a component of 250 to 900 Da with at least two rings, at most ten rotatable bonds and at least one nitrogen, and excluding cofactors and crystallization additives, leaves 5.9 percent. (b) The same filtering counted by protein, projected from 1,707 entries read in full across four accession shards, with a further gate on sequence identity to the donor structure and on local backbone RMSD at the transplant. About 1 in 8 proteins survive.

The accuracy of such transplants is difficult to establish. Their authors validated against experimental complexes of identical sequence, which existed for 0.24 percent of transplants, and reported that about 1% of that subset deviated by more than 2.5 Å in the local ligand environment. Their 2 confidence measures each correlate with that deviation at r = 0.51, so they are informative but not decisive, and no independent benchmark of transplanted ligand positions against experimentally determined ones has been published.

Redundancy is the more consequential obstacle. The proteins AlphaFill would contribute are reached through ligands the index already contains, because a transplant is by construction a igand already present in some solved structure. Across the qualifying ligands common to both resources, the number of targets attached to each would rise by roughly two orders of magnitude, so a retrieved neighbor would return a target pool larger than the set of sites the retrieval step exists to avoid docking into. The information added is that a ligand binds somewhere in a protein family, which a sequence search establishes more directly. The value of this method is in discriminating between candidate targets, and a source that returns whole protein families does not help with that.

The proteins most likely to benefit have no experimental complex and therefore no ground truth for evaluation. The index reported here is experimental throughout. Modeled complexes can be retrieved separately and presented alongside the experimental result.

### Software and data availability

The method is deployed at reversescreen.ai, which provides the fingerprints, the index and the code needed to screen in a local environment. PharmCast is at github.com/smuskal/pharmcast, and the model can be exercised on a single structure at pharmcast.ai. AutoDock Vina 1.2.7 is at github.com/ccsb-scripps/AutoDock-Vina and Meeko at github.com/forlilab/Meeko. All structures come from the RCSB Protein Data Bank, rcsb.org,^19,20^ and the component definitions from its chemical component dictionary. Target annotations are from ChEMBL.^21^

## Notes

https://reversescreen.ai

https://pharmcast.ai/

### Competing Interest Statement

The authors have declared no competing interest.

## References

1. Chen, Y. Z.; Zhi, D. G. Ligand-protein inverse docking and its potential use in the computer search of protein targets of a small molecule. Proteins 2001, 43, 217–226.

2. Li, H.; Gao, Z.; Kang, L.; Zhang, H.; Yang, K.; Yu, K.; Luo, X.; Zhu, W.; Chen, K.; Shen, J.; Wang, X.; Jiang, H. TarFisDock: a web server for identifying drug targets with docking approach. Nucleic Acids Res. 2006, 34, W219–W224.

3. Wang, J.-C.; Chu, P.-Y.; Chen, C.-M.; Lin, J.-H. idTarget: a web server for identifying protein targets of small chemical molecules with robust scoring functions and a divide-and-conquer docking approach. Nucleic Acids Res. 2012, 40, W393–W399.

4. Lapillo, M.; Tuccinardi, T.; Martinelli, A.; Macchia, M.; Giordano, A.; Poli, G. Extensive reliability evaluation of docking-based target-fishing strategies. Int. J. Mol. Sci. 2019, 20, 1023.

5. Keiser, M. J.; Roth, B. L.; Armbruster, B. N.; Ernsberger, P.; Irwin, J. J.; Shoichet, B. K. Relating protein pharmacology by ligand chemistry. Nat. Biotechnol. 2007, 25, 197–206.

6. Daina, A.; Zoete, V. Testing the predictive power of reverse screening to infer drug targets, with the help of machine learning. Commun. Chem. 2024, 7, 105.

7. McGregor, M. J.; Muskal, S. M. Pharmacophore fingerprinting. 1. Application to QSAR and focused library design. J. Chem. Inf. Comput. Sci. 1999, 39, 569–574.

8. McGregor, M. J.; Muskal, S. M. Pharmacophore fingerprinting. 2. Application to primary library design. J. Chem. Inf. Comput. Sci. 2000, 40, 117–125.

9. Muskal, S. M.; McGregor, M. J. PharmCast: rapid generation of three-dimensional pharmacophore fingerprints from two-dimensional structure without conformer generation. bioRxiv 2026, 10.64898/2026.09.02.748999.

10. Schmitt, S.; Kuhn, D.; Klebe, G. A new method to detect related function among proteins independent of sequence and fold homology. J. Mol. Biol. 2002, 323, 387–406.

11. Ehrt, C.; Brinkjost, T.; Koch, O. A benchmark driven guide to binding site comparison: an exhaustive evaluation using tailor-made data sets. PLoS Comput. Biol. 2018, 14, e1006483.

12. Trott, O.; Olson, A. J. AutoDock Vina: improving the speed and accuracy of docking with a new scoring function, efficient optimization, and multithreading. J. Comput. Chem. 2010, 31, 455–461.

13. Eberhardt, J.; Santos-Martins, D.; Tillack, A. F.; Forli, S. AutoDock Vina 1.2.0: new docking methods, expanded force field, and Python bindings. J. Chem. Inf. Model. 2021, 61, 3891–3898.

14. Riniker, S.; Landrum, G. A. Better informed distance geometry: using what we know to improve conformation generation. J. Chem. Inf. Model. 2015, 55, 2562–2574.

15. Howard, E. I.; Sanishvili, R.; Cachau, R. E.; Mitschler, A.; Chevrier, B.; Barth, P.; Lamour, V.; Van Zandt, M.; Sibley, E.; Bon, C.; Moras, D.; Schneider, T. R.; Joachimiak, A.; Podjarny, A. Ultrahigh resolution drug design I: details of interactions in human aldose reductase-inhibitor complex at 0.66 A. Proteins 2004, 55, 792–804.

16. Kawai, T.; Sun, B.; Yoshino, H.; Feng, D.; Suzuki, Y.; Fukazawa, M.; Nagao, S.; Wainscott, D. B.; Showalter, A. D.; Droz, B. A.;, et al. Structural basis for GLP-1 receptor activation by LY3502970, an orally active nonpeptide agonist. Proc. Natl. Acad. Sci. U. S. A. 2020, 117, 29959–29967.

17. Zhang, X.; Belousoff, M. J.; Zhao, P.; Kooistra, A. J.; Truong, T. T.; Ang, S. Y.; Underwood, C. R.; Egebjerg, T.; Senel, P.; Stewart, G. D.;, et al. Differential GLP-1R binding and activation by peptide and non-peptide agonists. Mol. Cell 2020, 80, 485–500.e7.

18. Jiang, J.; Jiang, L.; Maldonato, B. J.; Wang, Y.; Holderfield, M.; Aronchik, I.; Winters, I. P.; Salman, Z.; Blaj, C.; Menard, M.;, et al. Translational and therapeutic evaluation of RAS-GTP inhibition by RMC-6236 in RAS-driven cancers. Cancer Discov. 2024, 14, 994–1017.

19. Berman, H. M.; Westbrook, J.; Feng, Z.; Gilliland, G.; Bhat, T. N.; Weissig, H.; Shindyalov, I. N.; Bourne, P. E. The Protein Data Bank. Nucleic Acids Res. 2000, 28, 235–242.

20. Burley, S. K.; Bhikadiya, C.; Bi, C.;, et al. RCSB Protein Data Bank: powerful new tools for exploring 3D structures of biological macromolecules. Nucleic Acids Res. 2021, 49, D437–D451.

21. Zdrazil, B.; Felix, E.; Hunter, F.; Manners, E. J.; Blackshaw, J.; Corbett, S.; de Veij, M.; Ioannidis, H.; Lopez, D. M.; Mosquera, J. F.;, et al. The ChEMBL Database in 2023: a drug discovery platform spanning multiple bioactivity data types and time periods. Nucleic Acids Res. 2024, 52, D1180–D1192.

22. Hekkelman, M. L.; de Vries, I.; Joosten, R. P.; Perrakis, A. AlphaFill: enriching AlphaFold models with ligands and cofactors. Nat. Methods 2023, 20, 205–213. 10.1038/s41592-022-01685-y.

